# Individual variation in dispersal and fecundity increases rates of spatial spread

**DOI:** 10.1101/702092

**Authors:** Sebastian J. Schreiber, Noelle G. Beckman

**Affiliations:** Department of Evolution and Ecology and Center for Population Biology, University of California, Davis, California USA 95616; Department of Biology and Ecology Center, Utah State University, Logan, Utah 84322

## Abstract

Dispersal and fecundity are two fundamental traits underlying the spread of populations. Using integral difference equation models, we examine how individual variation in these fundamental traits and the heritability of these traits influence rates of spatial spread of populations along a one-dimensional transect. Using a mixture of analytic and numerical methods, we show that individual variation in dispersal rates increases spread rates and the more heritable this variation, the greater the increase. In contrast, individual variation in lifetime fecundity only increases spread rates when some of this variation is heritable. The highest increases in spread rates occurs when variation in dispersal positively covaries with fecundity. Our results highlight the importance of estimating individual variation in dispersal rates, dispersal syndromes in which fecundity and dispersal co-vary positively, and heritability of these traits to predict population rates of spatial spread.

---

Predicting the spatial spread of species over time is a central question in ecology [Hastings et al., 2005, Jongejans et al., 2008]. Mathematical models combining demography and dispersal have a long history of providing insights about the ecology and evolution of spatial spread [Skellam, 1951, Kot et al., 1996, Hastings et al., 2005, Beckman et al., in press]. These models have guided conservation and management decisions to control the spread of invasive species [e.g., Shea et al., 2010] and are used to make predictions about the persistence of species under shifting climates [e.g., Santini et al., 2016, Travis et al., 2011]. Traditionally, these models relied on mean estimates of dispersal and demographic rates. These rates, however, often exhibit substantial individual variation within populations [reviewed in Schupp et al., in revision]. As this individual variation is known to have important consequences for many ecological and evolutionary processes [Bolnick et al., 2011, Moran et al., 2016, Snell et al., 2019], it is natural to ask what effect do they have on rates of spatial spread.

In plants, variation in dispersal rates arises from intrinsic variation in trait expression among and within individuals and extrinsic variation based on the environmental context of the plant [Schupp et al., in revision, Saastamoinen et al., 2018]. Saastamoinen et al. [2018] found that while plants can have high levels of heritability in dispersal traits, there can be a wide range of heritability that depends on the specific trait measured and the environment in which it was measured. Theoretical studies have studied the effects of non-heritable and heritable variation in dispersal rates on spatial spread. Petrovskii and Morozov [2008] and Stover et al. [2014] found that non-heritable variation in dispersal rates, such as due to phenotypic plasticity in response to local environmental heterogeneity [Johnson et al., 2019], lead to fatter dispersal kernels and faster rates of spatial spread. Alternatively, theoretical studies accounting for only heritable variation found selection for increased dispersal rates on the edges of a species’ range resulting in accelerating rates of spatial spread [Travis and Dytham, 2002, Hughes et al., 2007, Phillips et al., 2008, Travis et al., 2009, Phillips et al., 2010, Bouin et al., 2012, Perkins et al., 2013]. Empirical studies of expanding plant populations have supported some of these theoretical predictions [Cwynar and MacDonald, 1987, Huang et al., 2015, Williams et al., 2016a, Tabassum and Leishman, 2018, 2019]. However, we still lack a full understanding of the relative contributions of heritable and non-heritable variation in dispersal rates on spread rates, and whether covariation among individuals in dispersal and demographic rates facilitate or constrain spread rates.

Rates of spatial spread are likely to depend on the covariance of dispersal with other traits under selection [Saastamoinen et al., 2018]. Within populations, higher fecundity in plants is expected to increase the distance seeds are dispersed [Clark et al., 1998a, Norghauer et al., 2011]. The number of fruit produced varies substantially among individuals within and across years in natural systems [e.g., Norghauer et al., 2011, Norghauer and Newbery, 2015] with moderate to high heritability found in crop systems [e.g., Jindal et al., 2010, Usman et al., 2014]. More generally, dispersal and life history traits may covary to produce integrated strategies known as dispersal syndromes [Ronce and Clobert, 2012] or dispersal may vary independently from other life history traits [Bonte and Dahirel, 2017]. Dispersal syndromes may arise due to a variety of proximate and ultimate causes [reviewed in Ronce and Clobert, 2012], including trade-offs in allocation, similar responses in expression to environmental conditions, genetic correlations among traits, joint selection on several traits, or selection on dispersal constrained by or constraining the evolution of other traits. Across species, Beckman et al. [2018] found species with fast life-history strategies dispersed their seeds further than species with slow life-history strategies. Within species, dispersal is predicted to be an independent axis of other life history traits [Bonte and Dahirel, 2017], although this is not well-studied in plants.

To better understand the simultaneous effects of heritable and non-heritable co-variation in dispersal and demographic rates on spatial spread, we introduce a new class of integral difference equation models. These spatially explicit models simultaneously account for individual variation in life-time fecundities and dispersal rates. This variation is allowed to be discrete or continuous, and heritable or non-heritable. Using this model, we explore the effects of variation in dispersal and demographic rates among individuals on the spread rate of populations, by first considering the separate effects of variation in dispersal and fecundity varying among individuals and then the joint effect of dispersal and fecundity covarying among individuals. Our mathematical analysis, buttressed by numerical simulations, highlights that individual variation in dispersal rates, generally, increases rates of spatial spread, while non-heritable variation in fecundity has no effect. In contrast, when individual variation in fecundity covaries positively with dispersal rates, it increases spread rates. Furthermore, heritability of either form of variation always increases spread rates.

## Model and Methods

Our models consider a population of plants living along a one-dimensional transect. Individuals vary in their production of seeds and the mean distance that a seed disperses. We consider two forms of the model: one with random transmission of individual traits and another allowing for non-random transmission of the traits. Both forms of the models are integro-difference equations that have been used extensively to model spatial spread [Kot et al., 1996, Neubert and Caswell, 2000]. For the model with non-random transmission, the population is structured by the trait in every spatial location. The changes in this local population structure are determined by a matrix model for discretely-structured traits and by an integral projection model for continuously-structured traits. For both types of models, we use the methods of Ellner and Schreiber [2012] to identify the asymptotic rates of spatial spread. Using these methods, we develop explicit formulas for how both forms of individual variation alter spatial rates of spread. As these formulas are derived in the limit of small indvidual variation, we also numerically investigate an empirically-based model to demonstrate that the insights from our formulas apply to larger amounts of individual variation.

### Models with random transmission

Let *n*_*t*_(*x*) denote the population density at location *x* in generation *t*. Under low-density conditions, individual plants produce *f* seeds during their life time. Each of these seeds disperse, on average, a distance of *ℓ* meters. We call this mean dispersal distance, the dispersal rate (i.e. the average number of meters a seed moves in a generation). The density of individuals with these characteristics equals *ρ*(*f, ℓ*). For seeds with a dispersal rate of 1 meter, let *k*_1_(*v*)*dv* be the infinitesimal probability that these seeds disperse from location *x* to location *x* + *v*. We assume that the dispersal kernel for a group of seeds with dispersal rate *ℓ* equals *k*_*R*_(*v*) = *k*_1_(*v/ℓ*)*/ℓ*i.e. the shape of the dispersal kernel is common to all seeds. The density of individuals with dispersal rate *ℓ* equals *ρ*_*L*_(*ℓ*) = *∫ρ*(*f, ℓ*) *df*. The population-level dispersal kernel corresponds to averaging dispersal kernels *k*_*R*_ across this individual variation (Fig. 1):

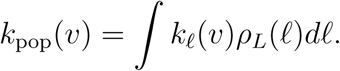

**Figure 1.**
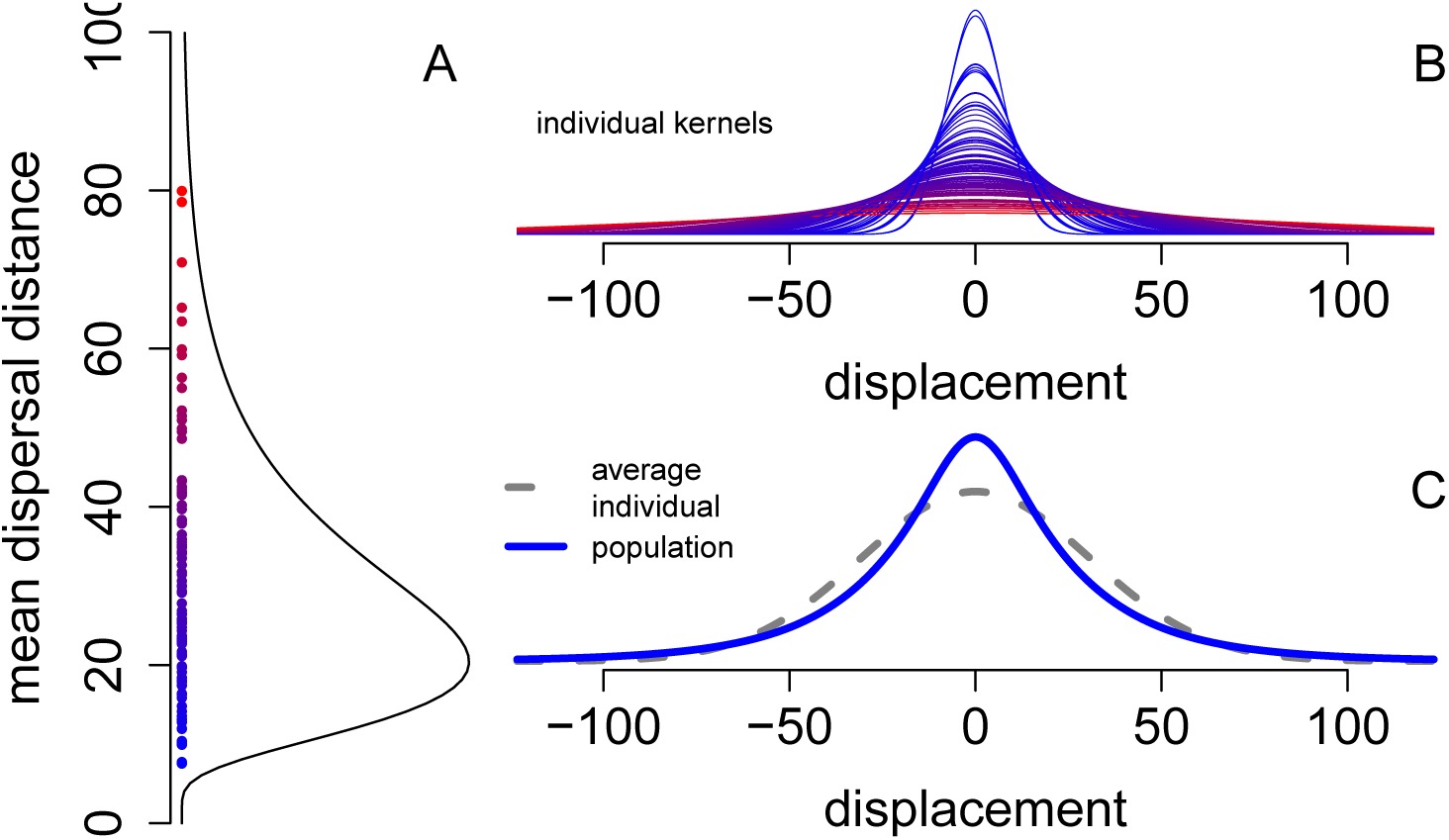
Individual variation in dispersal rates and the population-level dispersal kernel. In (A), variation among 100 maternal trees in their seed dispersal rates (mean dispersal distance). In (B), the Gaussian dispersal kernels of the 100 individuals from (A). In (C), the population-level dispersal kernel (i.e. the average of the kernels from (A)) in solid blue and the dispersal kernel of individuals with the average dispersal rate in dashed gray.

Petrovskii and Morozov [2008] call this population-level kernel a statistically structured dispersal model.

If *D*(*n*_*t*_(*y*)) corresponds to a density-dependent reduction in life-time fecundity at location *y*, then the spatial dynamics of the population is

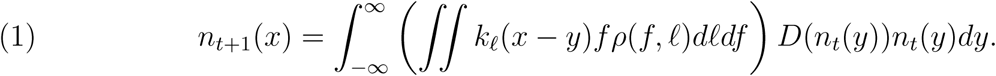

Without loss of generality, we assume that *D*(0) = 1. Furthermore, we assume that *D*(*n*) ≤ *D*(0) for all densities *n* ≥ 0 i.e. the lifetime fecundity of an individual is maximal at low densities. This assumption allows us to use the linearization principle for computing invasion speeds [Kot et al., 1996, Neubert and Caswell, 2000, Ellner and Schreiber, 2012].

While we have presented our model in equation (1) for continuously-structured traits, one can write a similar model for discretely-structured traits by replacing the double integral ∬ with a double sum Σ_*i*_ Σ_*j*_ and replacing the infinitesimal probabilities *ρ*(*f, ℓ*)*df dℓ* with discrete probabilities *ρ*(*f*_*i*_, *ℓ*_*j*_) for each of the traits. For example, the population-level dispersal kernel for discretely-structured population variation is 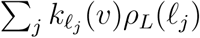 where *ρ*_*L*_(*ℓ*_*j*_) = Σ_*i*_ *ρ*(*f*_*i*_, *ℓ*_*j*_) is the marginal distribution of the individual dispersal rates (Fig. 1).

### Accounting for perfect transmission of traits

To account for seeds potentially inheriting their traits from their parents, we keep track of the density of individuals of a given trait combination at a given location. Specifically, let *n*_*t*_(*x*; *f, ℓ*) be the density of individuals of type *f, ℓ* at location *x* at time *t*. Let *ν* be the probability of perfect inheritance. When the trait isn’t perfectly transmitted, we assume that it is randomly transmitted with respect to the density *ρ*(*f, ℓ*). This model of inheritance provides a simple way to tune the heritability of traits from random transmission (*ν* = 0) to perfect transmission for all individuals (*ν* = 1). From a population genetics standpoint, this model corresponds to Turelli [1984]’s “House of Cards” model where mutations occurs with probability 1 − *ν* and the traits of the mutants are randomly drawn with respect to *ρ*(*f, ℓ*).

Under these assumptions, the model becomes

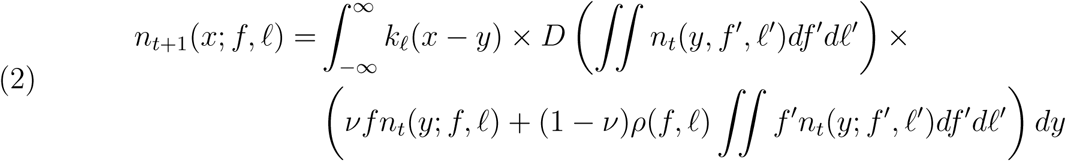

where *D* (*∬ n*_*t*_(*y, f*′, *ℓ*′)*df*′*dℓ*′) is the density-dependent reduction in fecundity at location *y* due to the total population density *∬ n*_*t*_(*y, f* ′, *ℓ*′)*df* ′*dℓ*′ at location *y*. For discretely-structured traits, we can use the same model structure by replacing the double integrals *∬ df dℓ* and *∬ df*′*dℓ*′ with a double sums Σ_*i*_ Σ_*j*_, and replacing the infinitesimal probabilities *ρ*(*f, ℓ*)*df dℓ* with discrete probabilities *ρ*(*f*_*i*_, *ℓ*_*j*_) for each of the traits.

### Analytic methods

To compute the asymptotic rates of spatial spread in both models, we make use of the linearization conjecture [Kot et al., 1996, Neubert and Caswell, 2000, Ellner and Schreiber, 2012] whose assumptions are satisfied whenever the base dispersal kernel *k*_1_(*v*) has exponentially bounded tails and the density *ρ*(*f, ℓ*) is compactly supported, that is, there exist *f*_min_< *f*_max_ and *ℓ*_min_ < *ℓ*_max_ such that 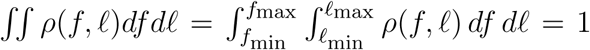. To use the linearization conjecture for the model with random transmission, we use the transform

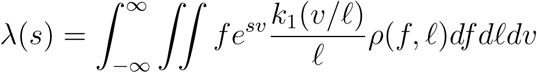

for the combined demography and dispersal kernel at low density. The linearization conjecture asserts that the asymptotic rate of spatial spread equals

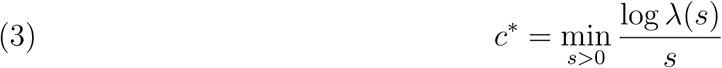

where the minimum is taken over values of s for which *λ*(*s*) is well-defined. Note, that equation (3) describes the spread rate on a generational time scale. To get a yearly rate of spread, we divide this generational rate of spread by the generation time in years.

For the model with perfect transmission, define the full demography and dispersal kernel *K* (*f, ℓ f*′, *ℓ, v*) by

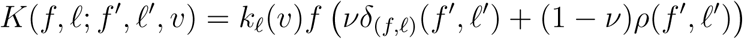

where *δ*_(*f,ℓ*)_(*f*′, *ℓ*′) is the Dirac delta function at (*f, ℓ*). Let *H*(*s*) be the operator that takes function of the form *n*(*f, ℓ*) to the function

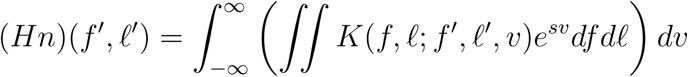

and let *λ*(*s*) be the dominant eigenvalue of *H*(*s*). Then the asymptotic rate of spatial spread is, once again, given by equation (3). When the individual variation is discretely structured, these formulas still apply but the double integrals *∬* need to be replaced with double sums Σ_*j*_ Σ_*j*_ and the density functions need to be replaced with probability distribution functions.

We use equation (3) in three ways. First, we approximate its solution for small variances. Namely, let *F* and *L* be random variables with joint density *ρ*(*f, ℓ*). Then, we can express these random variables in the form 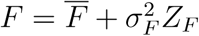 and 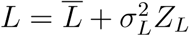, where 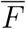 and 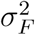 are the mean and variance of the fecundity, 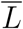 and 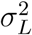 are the mean and variance of the mean dispersal distance *L*, and 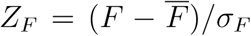 and 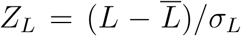 are random variables with a mean of 0 and variance of 1. In Appendix A, we derive approximations for the rates of spatial spread when 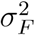 and 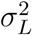 are sufficiently small. Second, to understand the effect of perfect transmission on rates of spatial spread, we use the reduction principle [Karlin, 1976, Altenberg and Feldman, 1987, Kirkland et al., 2006, Altenberg, 2012] in Appendix B to show that the rate of spatial spread increases with the probability *ν* of perfect transmission. Moreover, we derive an explicit approximation of the rate of spread for low levels of perfect transmission. While all of our analytical results apply both to continuous and discretely structured traits, we present the arguments in the Appendices for continuously structured traits. The same arguments apply to discretely structured traits by replacing integrals with sums.

Finally, we use equation (3) for our numerical calculations. The numerical calculations were based on empirical fits of dispersal data for the tree *Acer rubrum* [Clark et al., 1998b, Clark, 1998]. Clark et al. [1998b] collected data on seed rain over 5 years from 100 seed traps located within five 0.36-ha mapped tree stands in the southern Appalachians. When fit with a Gaussian dispersal kernel, the distance parameter *α* equals 30.8±3.80*SE* (average distance traveled is *α*Γ(1)*/*Γ(1*/*2) = 17.4). Clark [1998] estimated the net reproductive rate as 1, 325 and the generation time at *T* = 5.8 years. To get yearly rates of spread, we followed Clark [1998] and used *c/T*. The distribution of mean dispersal rates and fecundity were drawn from a hundred samples of a log-normal distribution with the variance and correlations reported in the figures.

## Results

Let 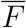 and 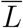 be the mean lifetime fecundity and mean dispersal rate of the population: 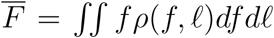 and 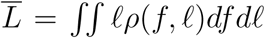. Let 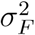 and 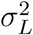 be the associated variances: 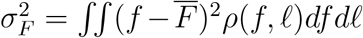 and 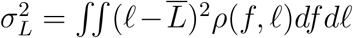. Let *r* be the correlation between the lifetime fecundity of individuals and the dispersal rates of their seeds: 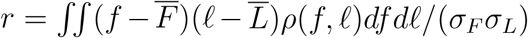. Let 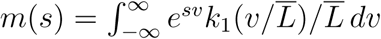 be the moment generating function for the dispersal kernel of the mean dispersal phenotype.

### Individual variation in dispersal rates

Using analytical approximations for small individual variation in dispersal rates, Appendix A demonstrates that randomly transmitted variation in dispersal rates increases the rate of spatial spread by a term proportional to the squared coefficient of variation in the mean dispersal distances. Specifically, for sufficiently small variance *σ*_*L*_, the increase is the spread rate equals

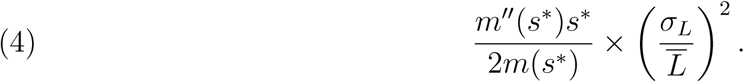

The proportionality constant 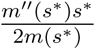 tends to increase with the variance in the base dispersal kernel; the greater this variance, the greater the increase in the rate of spread. Intuitively, if the base mode of dispersal has greater variation in distances traveled (e.g. the Laplacian kernel with a fatter tail versus the normal with a thinner tail), the greater the likelihood of individuals moving greater distances and it is these individuals that determine the rate of spatial spread.

When some of the variation in mean dispersal distances is perfectly transmitted to offspring, Appendix B shows that there always is an additional increase in the rate of spread. When the variation in dispersal rates is small and the probability of perfect transmission is small, this additional increase is proportional to the product of the coefficient of variation in the mean dispersal distance and the probability of perfect transmission. Specifically,

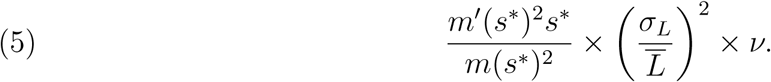

Consist with the analytical predications, numerical calculations for *Acer rubrum* based on equation (3) show that variation in dispersal rates and the heritability of this variation increase rates of spread (Figure 2A). However, at higher levels of variation, the approximation overestimates the spread rates (Figure 2B). None the less, the qualitative trends of variation in dispersal rates and perfect transmission of this variation increasing rates of spread still hold. Notably, even for moderate levels of variation and perfect transmission, individual variation in dispersal rates give substantial boosts to the predicted rate of spread for *Acer rubrum*. For example, a squared coefficient of variation of 0.5 more than doubles the rate of spatial spread (from approximately 20m/year to approximately 45m/year). If half of this variation is perfectly transmitted, then the spread rate nearly triples to 60m/year.

**Figure 2.**
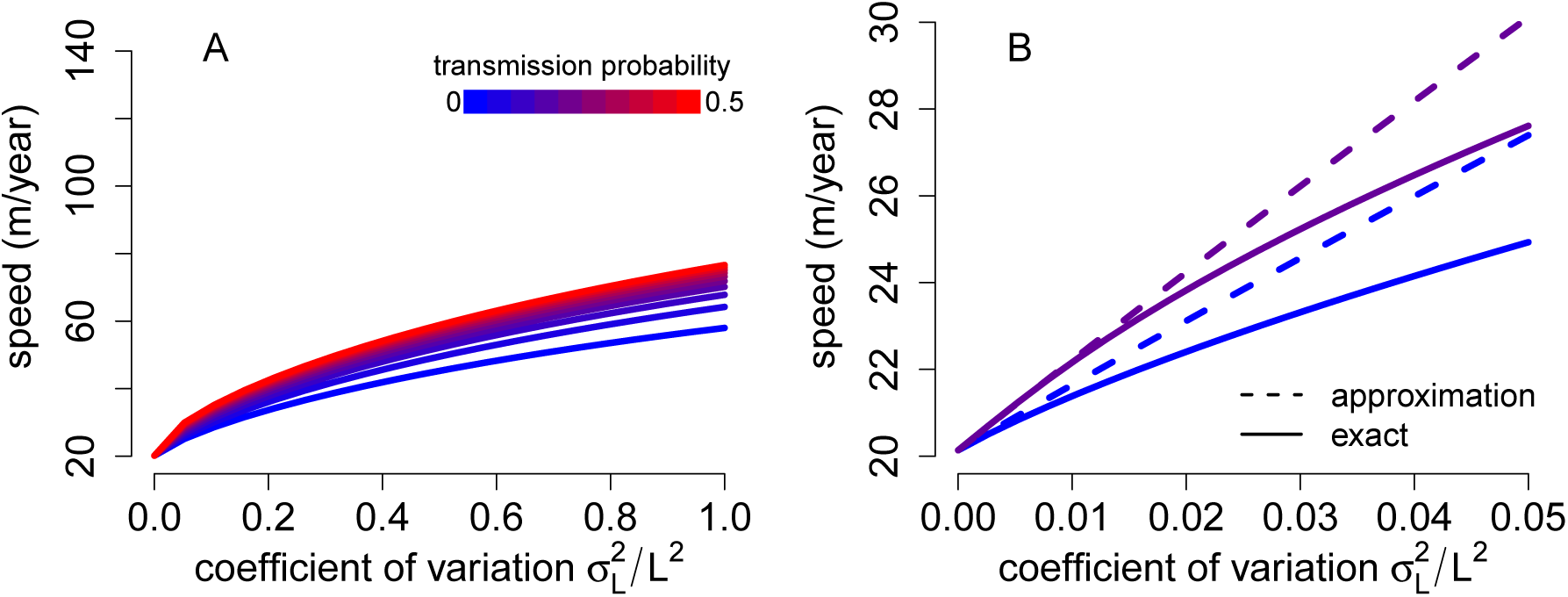
Individual variation in dispersal rates increases rate of spatial spread. In A, Rates of spatial spread for *Acer rubrum* (see *Methods*) are plotted against the coefficient of variation of the dispersal rate and for increasing probabilities of perfect transmission (from blue to red). In B, the analytical approximations (dashed lines) provide a good approximation to the exact invasion speeds (solid lines) for low variability and transmission probabilities. Higher levels of variation (A) have a decelerating effect on rates of spatial spread.

### Individual variation in fecundity

Randomly transmitted variation in fecundity has no effect on rates of spatial spread. However, when some of this variation is perfectly transmitted, Appendix B shows that there always is an increase in the spread rate. For low levels of individual variation in fecundity and perfect transmission, the invasion speed increases by a term proportional to the product of the squared coefficient of variation in fecundity and the probability of perfect transmission. Specifically,

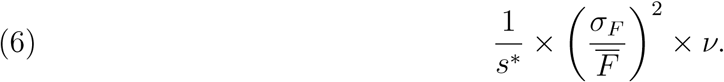

Figure 3 illustrates these effects numerically using equation (3) for the *Acer rubrum* model. In contrast to individual variation in dispersal rates, heritable variation in fecundity for this specific model has small effects on rates of spatial spread. For example, a coefficient of variation of 1 with a 50% chance of perfect transmission, speeds only increase approximately 9% for fecundity variation (Fig. 3A) in contrast to approximately 380% for dispersal variation (Fig. 2A). This relative small increase in the rate of spatial spread stems from the relatively small proportionality constant 1*/s*\* ≈ 8 in (6) compared to the proportionality constants in equation (4) with 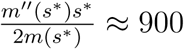 and equation (5) with 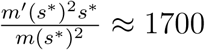.

**Figure 3.**
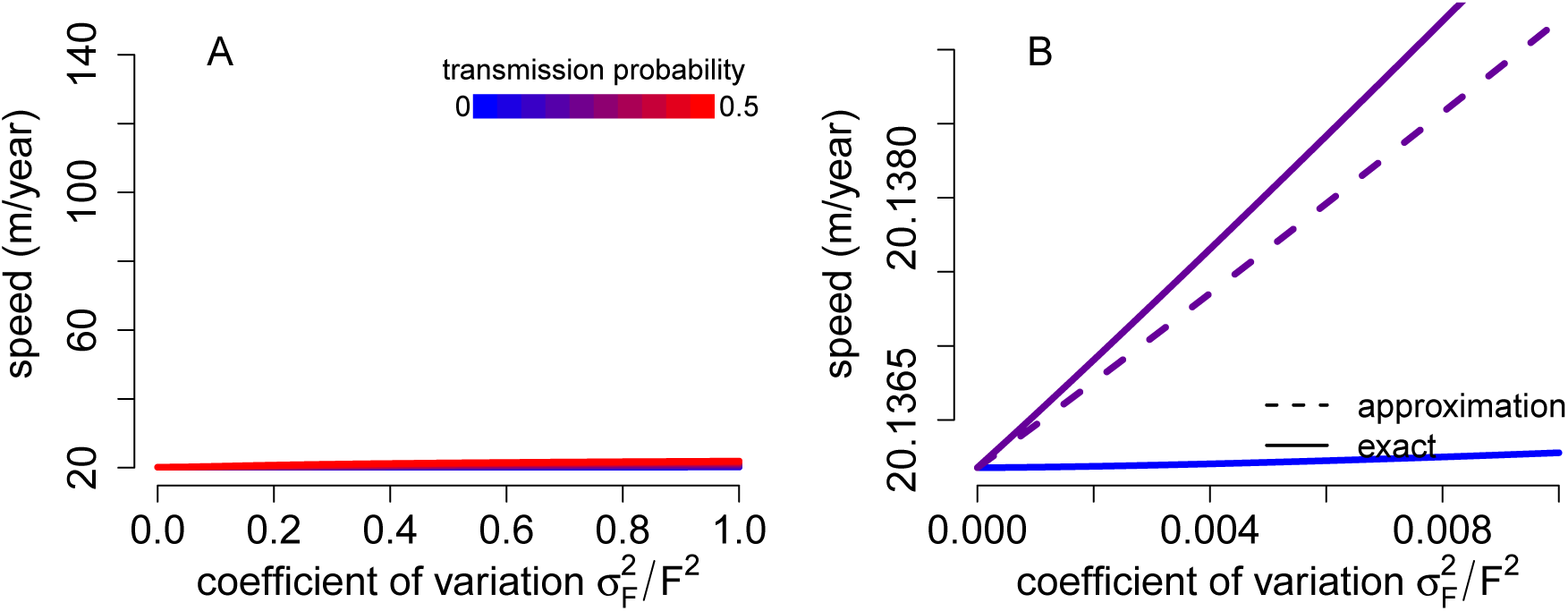
Individual variation in fecundity increases rate of spatial spread only when it is heritable. In A, invasion speeds for *Acer rubrum* (see *Methods*) are plotted against the coefficient of variation of fecundity and for increasing probabilities of perfect transmission (from blue to red). In B, for low variability and transmission probabilities, the analytical approximations (dashed lines) provide a good approximation to the exact invasion speeds (solid lines). Higher levels of variation (A) have nonlinear effects on these invasion speeds.

### Covariation in dispersal rates and fecundity

If lifetime fecundity of parents covary with dispersal rates of their seeds and this variation is randomly transmitted, then Appendix A shows that the spread rate increases by two terms: the amount due to dispersal variation alone in equation (4) plus an additional term proportional to the covariance of *L* and *F*:

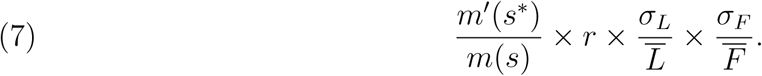

If this covariation is perfectly transmitted with probability *ν*, then Appendix B shows that there always is an additional increase to the spread rate. For low levels of individual variation and perfect transmission, this additional increase is proportional to the product of the covariance between fecundities and dispersal rates and the probability of perfect transmission:

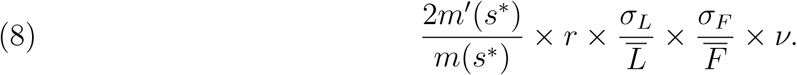

For the *Acer rubrum* model, Figure 4 illustrates the substantial increase due to this covariation: high positive correlation and heritability of individual variation in fecundity and dispersal rates (red curve in Fig. 4A) can lead to an eight-fold increase in the rate of spatial spread (approximately 160m/year) compared to the less than four fold increase (approximately 74m/year) due to uncorrelated variation in fecundity and dispersal rates (blue curve in Fig. 4A).

**Figure 4.**
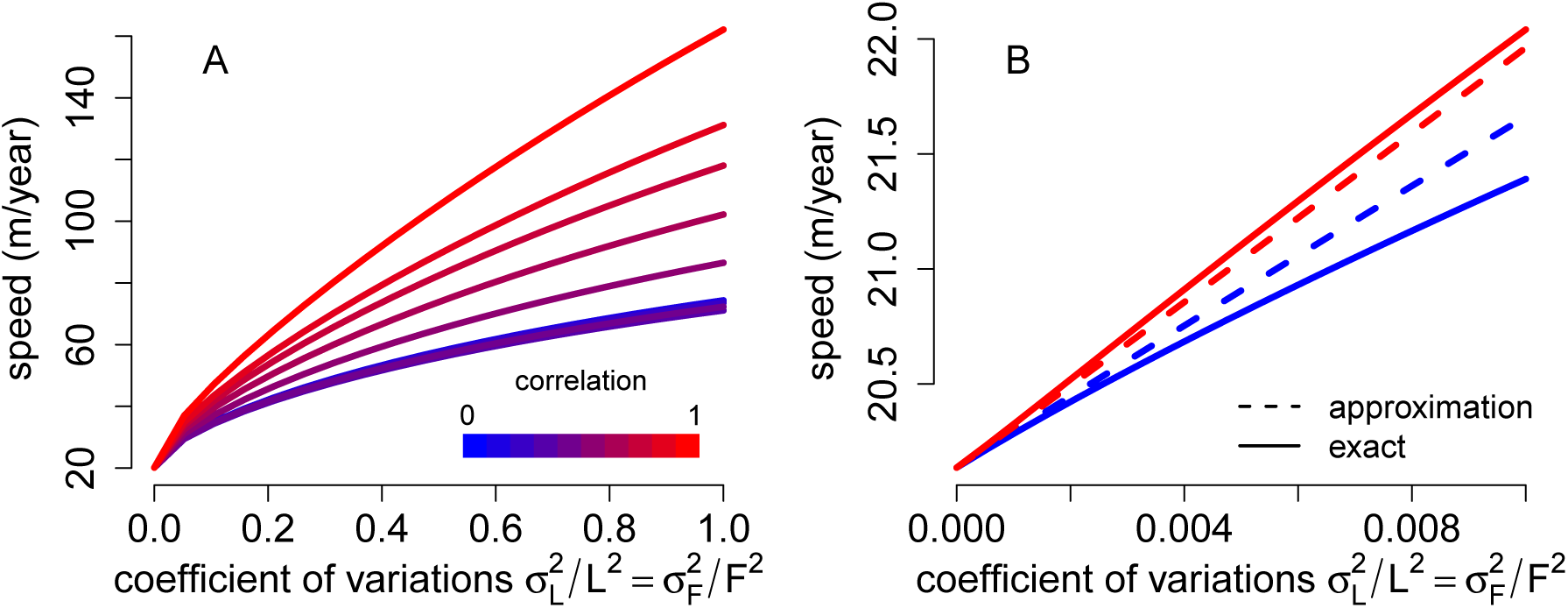
Covariation in fecundity and dispersal rates leads to faster rates of spatial spread. In A, spread rates for *Acer rubrum* (see *Methods*) are plotted against the coefficient of variations of fecundity and dispersal rates, and for increasing correlations between fecundity and dispersal rates (from blue to red). In B, for low variability, the analytical approximations (dashed lines) provide a good approximation to the exact invasion speeds (solid lines). Probability of perfect transmission is 0.5 in (A) and is 0.1 in (B).

## Discussion

Dispersal and fecundity are two fundamental traits underlying the spread of populations [Fisher, 1937, Skellam, 1951, Kot et al., 1996, Neubert and Caswell, 2000]. We show that inclusion of individual variation and covariation of these traits shifts predictions of population spread. Our results indicate that variation in dispersal increases spread rates of populations regardless of the mode of transmission, while variation in fecundity only increases spread rates when some of this variation is heritable. The highest increases in spread rates occurs when variation in dispersal positively covaries with fecundity. [Spread rates generally increase as heritability of dispersal rates and fecundity increase.] Although we focus on plants, our results are also applicable to animal systems

Our results are in-line with previous mathematical studies that show accelerated spread rates when individuals within the population vary in their dispersal ability [Bouin et al., 2012, Stover et al., 2014]. For gamma-distributed variation in dispersal rates and uniform distributions on two dispersal rates, Stover et al. [2014] showed that the moment generating functions of the population-level dispersal kernels increase with individual variation in dispersal rates and, thereby, increase spread rates. However, their numerical explorations found modest increases in spread rates when compared to our *Acer rubrum* example (e.g. about 20% in [Stover et al., 2014, Fig.3] versus 300% increase in spread rates for a squared coefficient of variation of 1). Our analytic approximation (see equation (4)) highlights that this type of difference stems from differences in mean dispersal rates. Specfically, the mean dispersal rate of *Acer rubrum* (30.8m/year) being greater than the base dispersal rate used by Stover et al. [2014] (1m/year). When intraspecific variability in dispersal rates is mostly heritable, Bouin et al. [2012] demonstrated that the spread rate is essentially determined by the genotypes with the highest dispersal rate being selected for at the edge of the spatial range, referred to as spatial sorting. Complementing this result, we used Karlin’s reduction principle [Karlin, 1976, Altenberg, 2012] to show that greater heritability leads to faster spread rates. Indeed, at low levels of heritability, equation (5) implies that the increase in spread rates is constrained by the coefficient of variation in the dispersal rates and the shape of population’s base dispersal kernel.

In contrast to individual variation in dispersal rates, we find that non-heritable variation in fecundity has no effect on rates of spatial spread. This outcome stems from (i) our analysis focusing on populations being sufficiently large that demographic stochasticity is negligible and (ii) the Laplace transform of the demography-dispersal kernel being a linear function of local demographic rates and a convex function of dispersal rates. As local demographic stochasticity slightly decreases spread rates [Snyder, 2003, Reluga, 2016] and individual variation in fecundity increases demographic stochasticity [Lloyd-Smith et al., 2005], it seems likely that demographic stochasticity coupled with individual variation in fecundity would decrease spread rates further. In contrast, we found that heritable variation in fecundity increases rates of spatial spread. In the extreme of this variation being perfectly transmitted from parents to offspring, we anticipate that spread rates are determined by selection for the most fecund individuals throughout the spatial range, unlike the spatial sorting mechanism for heritable variation in dispersal rates where selection only occurs at the edge of the spatial range Bouin et al. [2012].

We find the biggest effects of individual variation when dispersal rates and fecundity covary to form dispersal syndromes within species. Specifically, positive covariation of these traits, as has been found for some wind- and endozoochorous-dispersed seeds [reviewed in Schupp et al., in revision, Snell et al., 2019], always increases spread rates (e.g., more than doubling spread rates for *Acer rubrum*). Heritability of this covariation leads to greater increases of spatial spread. For example, our analysis implies that 50% heritability of this covariation can double the increase in spread rates (i.e. equations (7) and (8) are equal when *ν* = 0.5). In contrast, our analytic approximations in equations (7)-(8) imply that negative correlations between fecundity and dispersal rates lead to slower spread rates, but these rates are still higher than if there were no individual variation in fecundity or dispersal. Interestingly, Elliott and Cornell [2012] demonstrated that when there is trade-off between fecundity and dispersal (i.e. a negative correlation), polymorphisms of high and low fecundity individuals maintained by mutation lead to faster spread rates than the monomorphic spread rates. Whether these effects of covariation on spread rates are operating in natural systems remains to be seen.

Here we consider the influence of variation in dispersal, variation in fecundity, and their covariation on population spread rates under several simplifying assumptions. Understanding how relaxing these assumptions may alter these predictions provide many avenues for future research. Notably, we assumed the environment is spatially and temporally homogeneous. However, heterogeneous environments may alter these predictions. Heterogeneous environments can arise from natural disturbances, such as tree fall gaps, or through habitat loss and destruction due to human impacts. The latter tends to result in the fragmentation of the landscape into smaller, isolated fragments within a human-modified matrix. This fragmentation can alter rates of spatial spread [Shigesada et al., 1986, Kinezaki et al., 2010, Williams et al., 2016b, Crone et al., 2019]. For example, Shigesada et al. [1986] showed that habitat fragmentation slows down and, when sufficiently severe, halts spatial spread. Alternatively, temporal variation in fecundity and dispersal rates, respectively, slow down and speed up rates of spatial spread [Ellner and Schreiber, 2012]. To what extent heritable or non-heritable variation in dispersal rates and fecundity counter or amplify these effects of temporal and spatial heterogeneity remains to be understood. Furthermore, it would be useful to see how individual variation due to ontogenetic changes [Ellner et al., 2016] or genetics beyond the “house of cards” model [see, e.g., Johnson and Barton, 2005] influence our predictions about individual variation on spatial spread.

### Conclusion

Predictions of spread tend to rely on mean estimates of population parameters for dispersal and life history traits, but these may vary within a population and evolve through time. We found increased heritability in dispersal and fecundity increases spread rates compared to random transmission of traits, and if these are positively covarying to form dispersal syndromes within species, selection further facilitates increased spread rates. However, if dispersal and fecundity co-vary with other life history traits, selection for these traits may be constrained by or indirectly influence the evolution of other life-history traits, such as competitive ability or defense against natural enemies. The degree to which plant populations exhibit heritability of variation in dispersal or dispersal syndromes in which fecundity and dispersal co-vary positively is key to predicting the speed at which populations will track shifting habitats.

## Appendix A: Derivations for the random transmission model

In this Appendix derivations of the main analytic results are presented. As in the main text, let *F* and *L* be random variables with joint density function *ρ*(*f, ℓ*); more, generally *F* and *L* can be any mixture of a discrete and continuous distribution with finite moments. Then we can rewrite (1) as

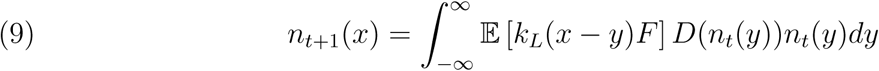

and *λ*(*s*) from the methods section in the main text can be rewritten as

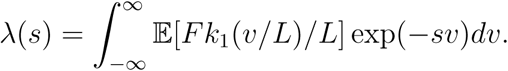

Provided that *F* and *L* have a finite variances, we can always write 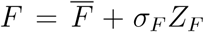 and 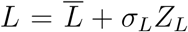 where *Z*_*F*_, *Z*_*L*_ are random variables with mean zero and variance 1, 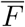 and 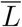 are the expected values of *F* and *L*, and 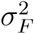 and 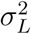 are the variances of *F* and *L*. Let *r* denote the correlation between *F* and *L*.

To derive the small variance approximations, we assume that there are positive constants *τ*_*F*_, *τ*_*L*_ such that *σ*_*F*_ = *ετ*_*F*_ and *σ*_*L*_ = *ετ*_*L*_ for small *ε* > 0. Ellner and Schreiber [2012] showed that

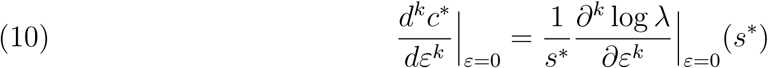

where *s*\* is such that *c*\* = *λ*(*s*\*)*/s*\* for *ε* = 0. By Tonelli’s theorem,

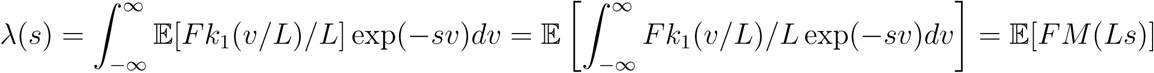

where 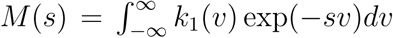. Differentiating with respect to *ε* and evaluating at zero yields

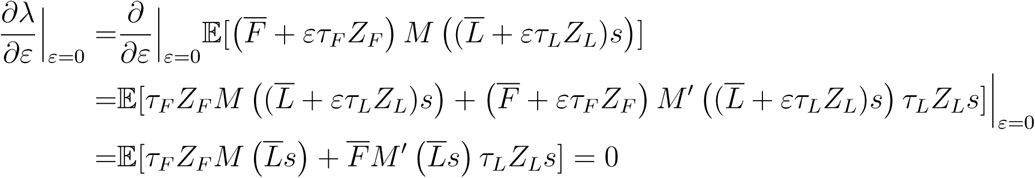

as 𝔼[*Z*_*F*_] = 𝔼[*Z*_*L*_] = 0. Hence, we get

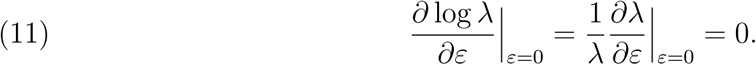

Differentiating a second time with respect to *ε* and evaluating at zero yields

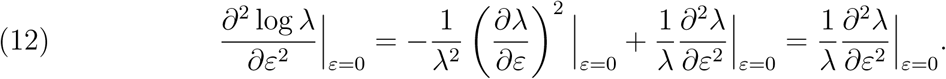

Computing the second derivative of *λ* yields

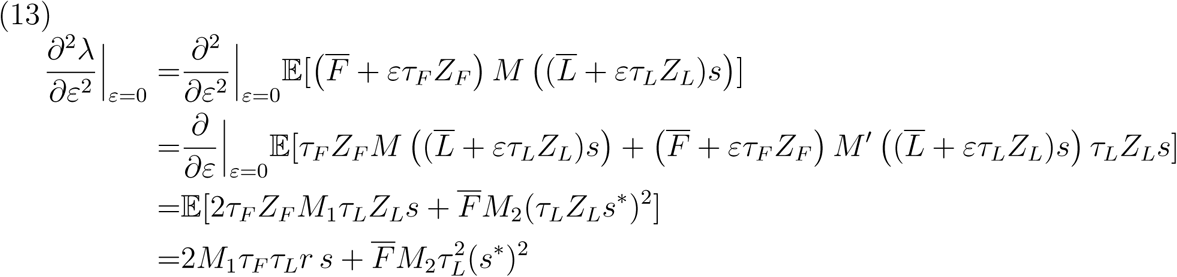

where 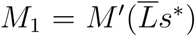 and 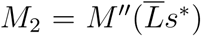. Recalling that *σ*_*F*_ = *ετ*_*F*_, *σ*_*L*_ = *ετ*_*L*_ and 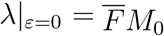 where 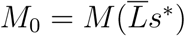, equations (10)–(13) give the second order approximation

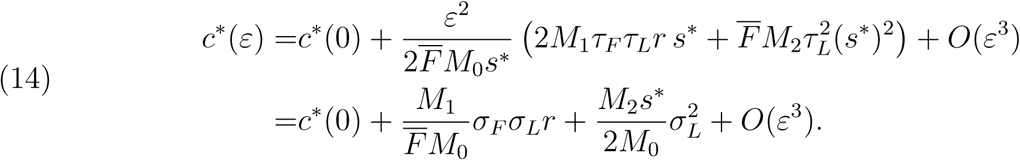

Defining 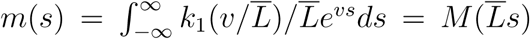, we get 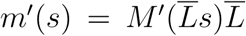 and 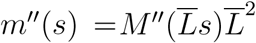. Hence, 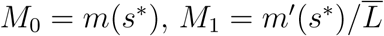 and 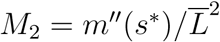 and

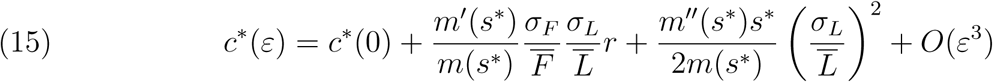

which gives equations (4) and (7) from the main text.

## Appendix B: Rates of spread for the model with perfect transmission

For the model with perfect transmission, recall that we have the demographic-dispersal kernel *K*_*ν*_(*f, ℓ*; *f* ′, *ℓ, v*) (now parameterized by *ν*) given by

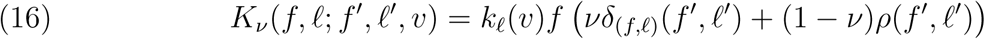

where *δ*_(*f,ℓ*)_(*f* ′, *ℓ*′) is the Dirac delta function based at the point (*f, ℓ*). Let *H*_*ν*_(*s*) be the operator that takes the function *n*(*f, ℓ*) to the function

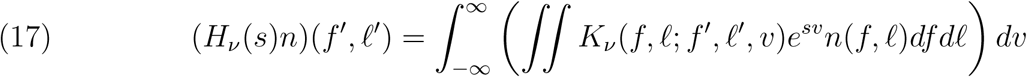

and let *λ*_*ν*_(*s*) be the dominant eigenvalue of *H*(*s*). The rate of spatial spread, as a function of *ν*, is given by 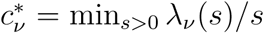. Let *s*\* be the value of *s* that gives the rate of spread for *ν* = 0 i.e. only random transmission.

Ellner and Schreiber [2012] showed that

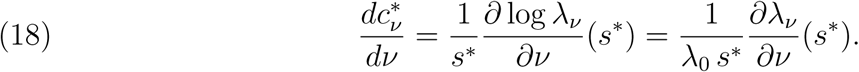

We will use the reduction principle [Karlin, 1976, Kirkland et al., 2006, Altenberg, 2012] to show that this derivative is always negative, i.e. the rate of spatial spread decreases with *ν*. Then we will compute this derivative at *ν* = 0 to get an approximation of the rate of spatial spread when *ν* is small.

We will show that the operator *H*_*ν*_(*s*) can be written in the form (*ν*Id + (1 *ν*)P) *D* where Id is the identity operator, *P* is a stochastic, positive operator, and *D* is a non-scalar multiplication operator. For operators of this form, Altenberg [2012, Theorem 6] proved that the dominant eigenvalue *λ*_*ν*_(*s*) is an increasing function of *ν*. Equivalently, *λ*_*ν*_(*s*) is a decreasing function of the probability 1 *ν* of mutation i.e. the reduction principle. Using the definition of *H*_*ν*_(*s*) and *K*_*ν*_ we have

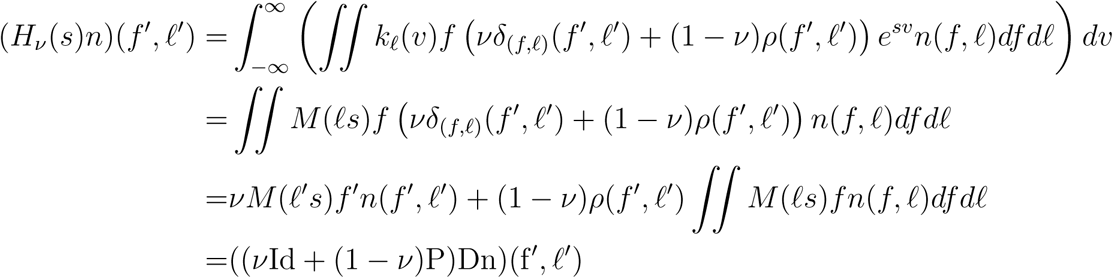

where *P* is the positive operator defined by (*Pn*)(*f* ′, *ℓ*′) = *ρ*(*f* ′, *ℓ*′) ∬ *n*(*f, ℓ*)*df dℓ* and *D* is the multiplication operator defined by (*Dn*)(*f* ′, *ℓ*′) = *M* (*ℓ*′*s*)*f* ′*n*(*ℓ*′, *f* ′). Hence, by the reduction principle *λ*_*ν*_(*s*) is an increasing function of *ν*. Equation (18) implies that 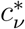 is an increasing function of *ν*.

To find the derivative in (18) at *ν* = 0, we need to find the dominant, left and right eigenfunctions of *H*_0_(*s*\*). The dominant, right eigenfunction *w* and eigenvalue *λ*_0_ must satisfy

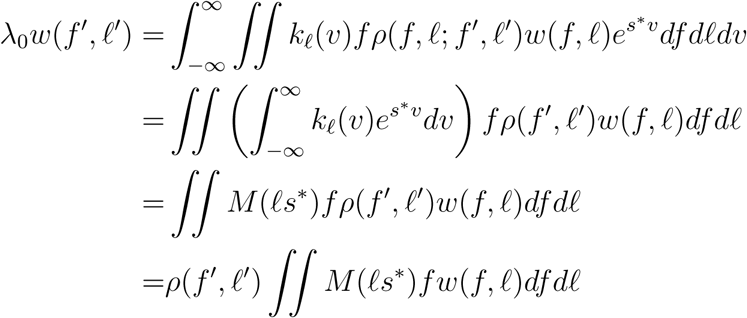

Hence, we get the unique, normalized eigenfunction is *w*(*f, ℓ*) = *ρ*(*f, ℓ*) and eigenvalue is *λ*_0_ = ∬ *M* (*ℓs*\*)*fρ*(*f, ℓ*)*df dℓ* = 𝔼[*FM* (*Ls*\*)] where (*F, L*) is the random vector with density function *ρ*(*f, ℓ*). The dominant, left eigenfunction *u*(*f, ℓ*) must satisfy

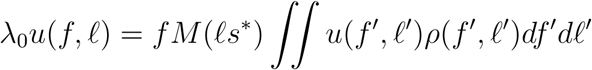

and therefore can be chosen to equal *u*(*f, ℓ*) = *f M* (*ℓs*\*). Hence, we get

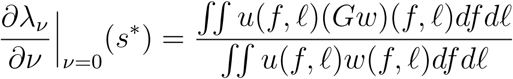

where *G* is the perturbation operator on functions *n*(*f, ℓ*) defined by

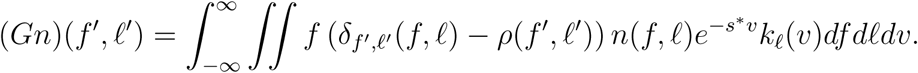

We have *∬u*(*f, ℓ*)*w*(*f, ℓ*)*df dℓ* = ∬*f M* (*ℓs*\*)*ρ*(*ℓ, f*)*dℓdf* = 𝔼[*FM* (*Ls*\*)] = *λ*_0_. Furthermore,

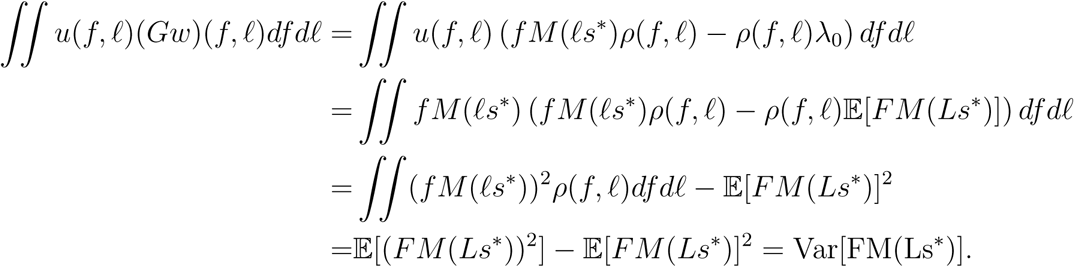

Thus, we get

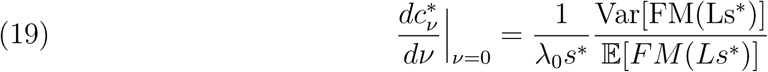

As in Appendix A, let 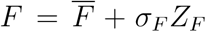 and 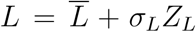 where *Z*_*F*_, *Z*_*L*_ are random variables with mean zero and variance 1, 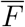 and 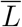 are the expected values of *F* and *L*, and 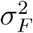 and 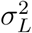 are the variances of *F* and *L*. Let *r* denote the correlation between *F* and *L*. To derive the small variance approximations, we assume that there are positive constants *τ*_*F*_, *τ*_*L*_ such that *σ*_*F*_ = *ετ*_*F*_ and *σ*_*L*_ = *ετ*_*L*_ for small *ε* > 0. With these assumptions, to get an approximation of Var[FM(Ls*)] for small *ε*, we need the following three approximations

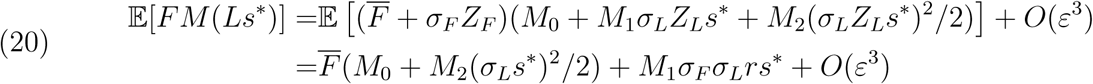

where 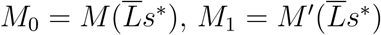 and 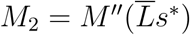, and

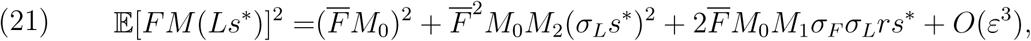

and

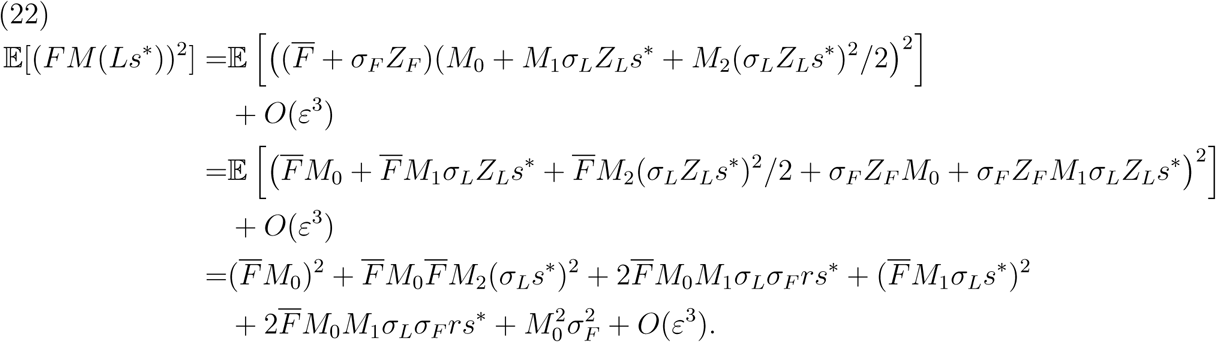

Taking the difference between (22) and (21) gives us

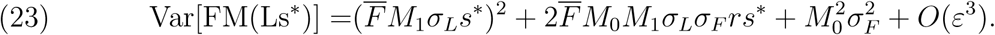

Thus, for sufficiently small *ν* and *ε*, equation (19) and 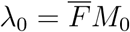 implies that

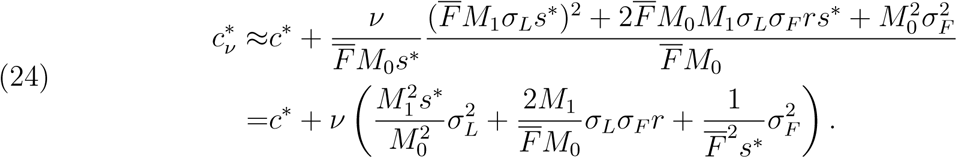

Defining 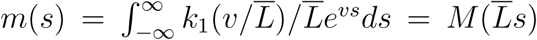, we get 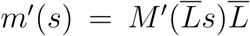 and 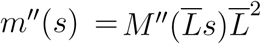. Hence, 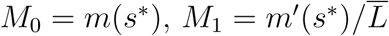 and 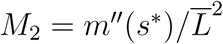 and

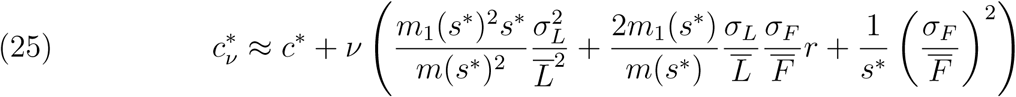

which gives equations (5), (6), and (8) from the main text.

We conclude by noting that provided they are bounded, all of the arguments here also apply when the joint distribution of *F* and *L* is any mixture of continuous and discrete distributions.

## References

L. Altenberg. Resolvent positive linear operators exhibit the reduction phenomenon. Proceedings of the National Academy of Sciences, 109:3705–3710, 2012.

L. Altenberg and M.W. Feldman. Selection, generalized transmission and the evolution of modifier genes. i. the reduction principle. Genetics, 117:559–572, 1987.

N. G. Beckman, J. M. Bullock, and R. Salguero-Gómez. High dispersal ability is related to fast life history strategies. Journal of Ecology, 106:1349–1362, 2018.

N. G. Beckman, C. E. Aslan, H. S. Rogers, O. Kogan, J. L. Bronstein, J. M. Bullock, F. Hartig, J. HilleRisLambers, Y. Zhou, D. Zurell, J. F. Brodie, E. M. Bruna, R. S. Cantrell, R. R. Decker, E. Effiom, E. C. Fricke, K. Gurski, A. Hastings, J. S. Johnson, B. A. Loiselle, M. N. Miriti, M. G. Neubert, L. Pejchar, J. R. Poulsen, G. Pufal, O. H. Razafindratsima, M. E. Sandor, K. Shea, S. Schreiber, E. W. Schupp, R. S. Snell, C. Strickland, and J. Zambrano. Advancing an interdisciplinary framework to study seed dispersal ecology. AoB Plants, in press.

D. I. Bolnick, P. Amarasekare, M. S. Araújo, R. Börger, J. M. Levine, M. Novak, V.H.W. Rudolf, S.J. Schreiber, M.C. Urban, and D.A. Vasseur. Why intraspecific trait variation matters in community ecology. Trends in Ecology & Evolution, 26:183–192, 2011.

Bonte M. Dahirel. Dispersal: a central and independent trait in life history. Oikos, 126:472–479, 2017.

Bouin, V. Calvez, N. Meunier, S. Mirrahimi, B. Perthame, G. Raoul, and R. Voituriez. Invasion fronts with variable motility: Phenotype selection, spatial sorting and wave acceleration. Comptes Rendus Mathematique, 350:761–766, 2012.

J. S. Clark, C. Fastie, G. Hurtt, S. T. Jackson, C. Johnson, G. A. King, M. Lewis, J. Lynch, S. Pacala, C. Prentice, E. W. Schupp, T. Webb, and P. Wyckoff. Reid’s paradox of rapid plant migration - dispersal theory and interpretation of paleoecological records. Bioscience, 48:13–24, 1998a.

J.S. Clark. Why trees migrate so fast: Confronting theory with dispersal biology and the paleorecord. The Americal Naturalist, 152:204–224, 1998.

J.S. Clark, E. Macklin, and L. Wood. Stages and spatial scales of recruitment limitation in southern appalachian forests. Ecological Monographs, 68:213–235, 1998b.

E.E. Crone, L.M. Brown, J.A. Hodgson, F. Lutscher, and C.B. Schultz. Faster movement in nonhabitat matrix promotes range shifts in heterogeneous landscapes. Ecology, page e02701, may 2019. doi: 10.1002/ecy.2701. URL https://doi.org/10.1002%2Fecy.2701.

L. C. Cwynar and G. M. MacDonald. Geographical variation of lodgepole pine in relation to population history. The Americal Naturalist, 129:463–469, 1987.

E.C. Elliott and S.J. Cornell. Dispersal polymorphism and the speed of biological invasions. PloS one, page e40496, 2012.

S.P. Ellner and S.J. Schreiber. Temporally variable dispersal and demography can accelerate the spread of invading species. Theoretical Population Biology, 82:283–298, 2012.

S.P. Ellner, D.Z. Childs, and M. Rees. Data-driven modelling of structured populations. Springer, 2016.

R. A. Fisher. The wave of advance of advantageous genes. Annals of Eugenics, 7:355–369, 1937.

A. Hastings, K. Cuddington, K. F. Davies, C. J. Dugaw, S. Elmendorf, A. Freestone, S. Harrison, M. Holland, J. Lambrinos, U. Malvadkar, B. A. Melbourne, K. Moore, C. Taylor, and D. Thomson. The spatial spread of invasions: new developments in theory and evidence. Ecology Letters, 8:91–101, 2005.

F. Huang, S. Peng, B. Chen, H. Liao, Q. Huang, Z. Lin, and G. Liu. Rapid evolution of dispersal-related traits during range expansion of an invasive vine *Mikania micrantha*. Oikos, 124:1023–1030, 2015.

C.L. Hughes, C. Dytham, and J.K. Hill. Modelling and analysing evolution of dispersal in populations at expanding range boundaries. Ecological Entomology, 32:437–445, 2007.

S. K. Jindal, D. Arora, and T. R. Ghai. Variability studies for yield and its contributing traits in okra. Electronic Journal of Plant Breeding, 1:1495–1499, 2010.

J. S. Johnson, R. S. Cantrell, C. Cosner, F. Hartig, A. Hastings, H. S. Rogers, E. W. Schupp, K. Shea, B. J. Teller, X. Yu, D. Zurell, and G. Pufal. Rapid changes in seed dispersal traits may modify plant responses to global change. AoB Plants, page plz020, 2019.

T. Johnson and N. Barton. Theoretical models of selection and mutation on quantitative traits. Philosophical Transactions of the Royal Society B: Biological Sciences, 360:1411–1425, 2005.

E. Jongejans, O. Skarpaas, and K. Shea. Dispersal, demography and spatial population models for conservation and control management. Perspectives in Plant Ecology Evolution and Systematics, 9:153–170, 2008.

S Karlin. Population subdivision and selection migration interaction. In Population genetics and ecology, pages 617–657. Academic Press New York, 1976.

Noriko Kinezaki, Kohkichi Kawasaki, and Nanako Shigesada. The effect of the spatial configuration of habitat fragmentation on invasive spread. Theoretical Population Biology, 78 (4):298–308, dec 2010. doi: 10.1016/j.tpb.2010.09.002. URL https://doi.org/10.1016%2Fj.tpb.2010.09.002.

S. Kirkland, C.K. Li, and S.J. Schreiber. On the evolution of dispersal in patchy landscapes. SIAM Journal on Applied Mathematics, 66(4):1366–1382, 2006.

M. Kot, M. A. Lewis, and P. van den Driessche. Dispersal data and the spread of invading organisms. Ecology, 77:2027–2042, 1996.

J.O. Lloyd-Smith, S.J. Schreiber, P.E. Kopp, and W.M. Getz. Superspreading and the effect of individual variation on disease emergence. Nature, 438:355–9, 2005.

E. V. Moran, F. Hartig, and D. M. Bell. Intraspecific trait variation across scales: implications for understanding global change responses. Global Change Biology, 22:137–150, 2016.

M.G. Neubert and H. Caswell. Demography and dispersal: Calculation and sensitivity analysis of invasion speed for structured populations. Ecology, 81:1613–1628, 2000.

J. M. Norghauer and D. M. Newbery. Tree size and fecundity influence ballistic seed dispersal of two dominant mast-fruiting species in a tropical rain forest. Forest Ecology and Management, 338:100–113, 2015.

J. M. Norghauer, C. A. Nock, and J. Grogan. The importance of tree size and fecundity for wind dispersal of big-leaf mahogany. PloS One, 6:e17488, 2011.

T. A. Perkins, B. L. Phillips, M. L. Baskett, and A. Hastings. Evolution of dispersal and life history interact to drive accelerating spread of an invasive species. Ecology Letters, 16: 1079–1087, 2013.

S. Petrovskii and A. Morozov. Dispersal in a statistically structured population: fat tails revisited. The American Naturalist, 173:278–289, 2008.

B. L. Phillips, G. P. Brown, and R. Shine. Life-history evolution in range-shifting populations. Ecology, 91:1617–1627, 2010.

B.L. Phillips, G.P. Brown, J.M.J. Travis, and R. Shine. Reid’s paradox revisited: the evolution of dispersal kernels during range expansion. American Naturalist, 172:S34–S48, 2008.

T.C. Reluga. The importance of being atomic: Ecological invasions as random walks instead of waves. Theoretical Population Biology, 112:157–169, 2016.

Ophelie Ronce and Jean Clobert. Dispersal Syndromes, book section 10, pages 119–138. Oxford University of Press, United Kingdom, 2012.

M. Saastamoinen, G. Bocedi, J. Cote, D. Legrand, F. Guillaume, C. W. Wheat, E.A. Fronhofer, C. Garcia, R. Henry, A. Husby, M. Baguette, D. Bonte, A. Coulon, H. Kokko, E. Matthysen, K. Niitepõld, E. Nonaka, V. M. Stevens, J. M. J. Travis, K. Donohue, J. M. Bullock, and M. del Mar Delgado. Genetics of dispersal. Biological Reviews, 93:574–599, 2018.

L. Santini, T. Cornulier, J.M. Bullock, S.C.F. Palmer, S.M. White, G. Bocedi, C. Rondinini, and J.M.J. Travis. Modelling spread rate in terrestrial mammals and the ability to track a shifting climate: a trait space approach. Global Change Biology, 22:2415–2424, 2016.

E. W. Schupp, R. Zwolak, Jones L. R, R. S. Snell, N. G. Beckman, C. Aslan, B. R. Cavazos, E. Effiom, E. C. Fricke, F. Montaño-Centellas, J. Poulsen, O. H. Razafindratsima, M. E. Sandor, and K. Shea. Intrinsic and extrinsic drivers of intraspecific variation in seed dispersal are diverse and pervasive. AoB Plants, in revision.

K. Shea, E. Jongejans, O. Skarpaas, D. Kelly, and A. W. Sheppard. Optimal management strategies to control local population growth or population spread may not be the same. Ecological Applications, 20:1148–1161, 2010.

N. Shigesada, K. Kawasaki, and E. Teramoto. Traveling periodic waves in heterogeneous environments. Theoretical Population Biology, 30(1):143–160, 8 1986. doi: 10.1016/0040-5809(86)90029-8. URL http://dx.doi.org/10.1016/0040-5809(86)90029-8.

J. G. Skellam. Random dispersal in theoretical populations. Biometrika, 38:196–218, 1951.

R.S. Snell, N.G. Beckman, E. Fricke, B.A. Loiselle, C.S. Carvalho, L.R. Jones, N.I. Lichti, N. Lustenhouwer, S. Schreiber, C. Strickland, L.L. Sullivan, B.R. Cavazos, I. Giladi, A. Hastings, K. Holbrook, E. Jongejans, O. Kogan, F. Montaño-Centellas, J. Rudolph, H. S Rogers, R. Zwolak, and E. Schupp. Consequences of intraspecific variation in seed dispersal for plant demography, communities, evolution, and global change. AoB PLANTS, 03 2019.

R.E. Snyder. How demographic stochasticity can slow biological invasions. Ecology, pages 1333–1339, 2003.

J.P. Stover, B.E. Kendall, and R.M. Nisbet. Consequences of dispersal heterogeneity for population spread and persistence. Bulletin of Mathematical Biology, 76:2681–2710, 2014.

S. Tabassum and M. R. Leishman. Have your cake and eat it too: greater dispersal ability and faster germination towards range edges of an invasive plant species in eastern australia. Biological Invasions, 20:1199–1210, 2018.

S. Tabassum and M. R. Leishman. It doesn’t take two to tango: increased capacity for self-fertilization towards range edges of two coastal invasive plant species in eastern australia. Biological Invasions, 21:1–13, 2019.

J. M. J. Travis and C. Dytham. Dispersal evolution during invasions. Evolutionary Ecology Research, 4:1119–1129, 2002.

J. M. J. Travis, K. Mustin, T. G. Benton, and C. Dytham. Accelerating invasion rates result from the evolution of density-dependent dispersal. Journal of Theoretical Biology, 259: 151–158, 2009.

J. M. J. Travis, Catriona M. Harris, K. J. Park, and J. M. Bullock. Improving prediction and management of range expansions by combining analytical and individual-based modelling approaches. Methods in Ecology and Evolution, 2:477–488, 2011.

M. Turelli. Heritable genetic variation via mutation-selection balance: Lerch’s zeta meets the abdominal bristle. Theoretical Population Biology, 25:138–193, 1984.

M. G. Usman, M. Y. Rafii, M. R. Ismail, M. A. Malek, and M. Abdul Latif. Heritability and genetic advance among chili pepper genotypes for heat tolerance and morphophysiological characteristics. The Scientific World Journal, 2014:14, 2014.

J. L. Williams, B. E. Kendall, and J. M. Levine. Rapid evolution accelerates plant population spread in fragmented experimental landscapes. Science, 353:482–485, 2016a.

J. L. Williams, R. E. Snyder, and J. M. Levine. The influence of evolution on population spread through patchy landscapes. American Naturalist, 188:15–26, 2016b.

